# RNA polymerase II O-GlcNAcylation promotes nuclear entry to drive transcription

**DOI:** 10.1101/2025.03.18.643968

**Authors:** Satu Pallasaho, Xun Lu, Aishwarya Gondane, Harri M Itkonen

## Abstract

O-GlcNAcylation of RNA Polymerase II (RNA Pol II) was described in 1993 and here we wanted to probe the functional significance of this modification. O-GlcNAc transferase (OGT) is the sole enzyme performing nucleo-cytoplasmic O-GlcNAcylation. The role of RNA Pol II O-GlcNAcylation has remained enigmatic partly due to the lack of reagents selectively recognizing the O-GlcNAcylated polymerase. We developed a novel approach, wheat germ agglutinin (WGA) lectin-based enrichment for O-GlcNAcylated proteins followed by traditional ChIP for RNA Pol II, to identify chromatin-binding sites of the O-GlcNAcylated RNA Pol II. We termed this technique as wChIP and used it to demonstrate that the O-GlcNAcylated RNA Pol II is enriched to the promoters of highly expressed genes. RNA Pol II has a repetitive c-terminal domain (CTD) of YSPTSPS, which is heavily post-translationally modified. Using structural modeling and WGA pulldown-experiments, we identify Ser-5 as a CTD site that is dynamically modified by OGT. Mass spectrometry-based detection of proteins that recognize CTD that is O-GlcNAcylated on Ser-5 identified nuclear transport protein, karyopherin subunit beta 1 (KPNB1), as a reader protein for this modification. Functionally, high activity of both OGT and KPNB1 is required for the efficient entry of RNA Pol II to nucleus and chromatin. OGT and KPNB1 are significantly co-expressed in prostate cancer tissue, and their combined inhibition is more toxic to prostate cancer cells than normal prostate cells. We propose that RNA Pol II O-GlcNAcylation promotes entry to nucleus, and this could integrate metabolic information to regulate global transcription.

## Introduction

RNA polymerase II (RNA Pol II) transcribes the protein-encoding part of the genome. The carboxy-terminal domain (CTD) of RNA Pol II has an important role in transcription regulation. Phosphorylation of the RNA Pol II CTD by cyclin-dependent kinases (CDKs) regulates transcription in all eukaryotes [1]. CDK7 phosphorylates the CTD at serine 5 to promote transcription initiation while CDK9-mediated phosphorylation at serine 2 promotes release from promoter-proximal pausing. CDK12 and CDK13 maintain the serine 2 phosphorylation status on particularly the long genes to sustain transcription elongation.

In addition to phosphorylation, the RNA Pol II CTD is also modified by O-GlcNAc transferase (OGT) [2-4]. OGT catalyzes the addition of a single N-Acetylglucosamine (GlcNAc) unit to the target protein serine or threonine residues in a process referred to as O-GlcNAcylation [5]. This modification can be removed by O-GlcNAcase (OGA) [6]. The OGT-OGA enzyme pair is responsible for all nucleocytoplasmic O-GlcNAcylation [7, 8]. RNA Pol II CTD contains four possible O-GlcNAcylation sites: serine 2, threonine 4, serine 5, and serine 7 [2-4]. RNA Pol II O-GlcNAcylation is important for promoter entry in partially purified transcription assays [3, 4]. However, it is not known where the O-GlcNAcylated polymerase is located across the genome and what proteins bind to the modified polymerase.

Co-targeting of OGT and transcriptional CDKs is toxic to prostate cancer cells [9-12], and OGT and transcriptional kinases may compete for CTD as substrate. Indeed, *in vitro* assays have shown that O-GlcNAcylation and phosphorylation are mutually exclusive on synthetic CTD peptide [13]. Both OGT and O-GlcNAcylation are enriched on the transcriptionally active chromatin [14-16]. However, the lack of reagents selectively recognizing O-GlcNAcylated RNA Pol II has made it difficult to study how O-GlcNAcylation affects the polymerase function.

Increased OGT expression is a feature of practically all cancers, including prostate cancer which is the second most common cancer among men [17-21]. Prostate cancer tumour growth is regulated by transcriptional programs driven by a hyper-activated transcription factor, androgen receptor [22]. Early-stage prostate cancer is successfully treated with androgen-deprivation therapy. However, despite the initial success of this treatment, the disease often progresses to a treatment-resistant form referred to as castration-resistant prostate cancer (CRPC) that currently lacks curative treatment options. OGT levels and O-GlcNAcylation are elevated in aggressive prostate cancer, and OGT inhibition decreases proliferation of the prostate cancer cells [19, 23, 24]. Additionally, we have previously shown that androgen receptor drives metabolic processes that enhance OGT activity [19, 25]. Prostate cancer therefore represents an ideal model system to explore the significance of RNA Pol II O-GlcNAcylation.

Here, we develop a tool to map the chromatin localization of O-GlcNAcylated RNA Pol II, lectin-based WGA re-ChIP (wChIP) and show that O-GlcNAcylated RNA Pol II is enriched on the promoters of highly transcribed genes. We show that inhibition of CDK7 stimulates RNA Pol II O-GlcNAcylation and identify O-GlcNAcylation of CTD Ser-5 as regulator of nuclear entry. In more detail, nuclear transport protein, karyopherin subunit beta 1 (KPNB1), binds to the O-GlcNAcylated Ser-5 and co-targeting of OGT and KPNB1 decreases both nuclear and chromatin-bound RNA Pol II. Finally, we show that cancer cells benefit from high activity of both OGT and KPNB1: the two are co-expressed in patient tumours and their co-targeting is toxic to prostate cancer cells. We propose that O-GlcNAcylation promotes RNA Pol II nuclear entry and cancer cells highjack this process to sustain high levels of transcription.

## Results and discussion

### Common chromatin O-GlcNAcylation sites across cell lines are enriched at promoters

We hypothesized that chromatin O-GlcNAcylation is located in specific regions of the genome in most cell types. To probe this, we identified the pan-cell line sites of the O-GlcNAcylated chromatin using seven previously published datasets enriching for O-GlcNAcylated chromatin (colon cancer (HT-2, GSE93656) [26], human embryonic kidney (HEK293, GSE51673) [27], Burkitt lymphoma (BJAB, GSE86154) [28], human embryonic kidney (HEK293, GSE36620) [14], androgen sensitive-positive prostate cancer (LNCaP, GSE121474) [15], androgen insensitive prostate cancer (PC3, GSE121474) [15]) and breast cancer (MCF-7, GSE141698) [16]). We identified 92 O-GlcNAc chromatin loci common for all the seven datasets (**Fig. 1A**). Practically all of these sites were located on promoter regions (98%) (**Fig. 1B**). In contrast, when evaluating the O-GlcNAc site distribution in the individual datasets, the enrichment for the promoters was between 22 to 88% (**Fig. 1C**). The fact that all the assessed datasets identify a relatively small number of sites that are highly enriched on the promoters implies that chromatin O-GlcNAcylation regulates transcription of a specific set of genes in all cells and may also regulate a distinct aspect of transcription. We assessed if the pan-cell line O-GlcNAc-regulated genes are related to a specific biological process. However, we were unable to identify any significantly enriched process (**Suppl. Fig. 1**).

**Figure 1.**
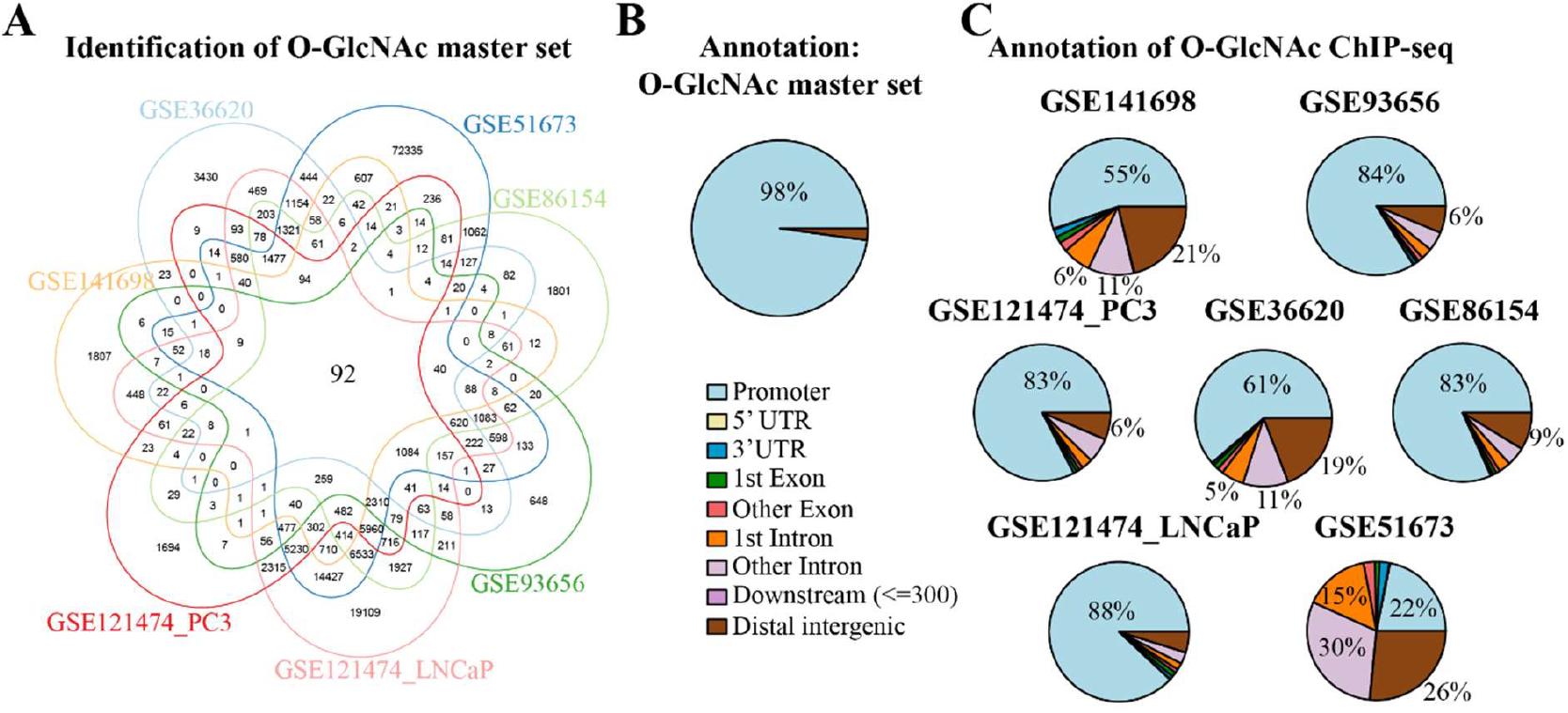
Chromatin O-GlcNAcylation is found predominantly at promoters. **A)** Venn diagram presenting the overlaps between the O-GlcNAcylated chromatin sites identified from seven datasets collected from six different cell lines: colon cancer cells (HT-2, GSE93656) [26], human embryonic kidney cells (HEK293, GSE51673) [27], Burkitt lymphoma cells (BJAB, GSE86154) [28], breast cancer cells (MCF-7, GSE141698) [16], HEK293, GSE36620) [14] and two prostate cancer cell lines (LNCaP and PC3) [15]. Genomic intervals at which the overlaps occur were counted using Bedtools multiinter [57]. **B-C)** Annotation of the O-GlcNAc sites of each of the seven datasets and the sites overlapped in all the datasets. Annotation was performed with ChIPseeker [64]. Percentage of annotation is indicated when value is higher than 5%.

We have so far shown that O-GlcNAcylated sites are enriched on the promoters. An obvious limitation of our approach is that we do not know which actual protein(s) is O-GlcNAcylated. The common factor present in the promoters of all cells is RNA polymerase II, the core component of the transcription machinery. We moved on to probe how O-GlcNAcylation regulates the chromatin localization of RNA Pol II.

### Development and validation of a re-ChIP strategy to enrich O-GlcNAcylated RNA Pol II

We hypothesized that O-GlcNAcylation regulates RNA Pol II localization on chromatin. Currently, there is no antibody that would be specific against the O-GlcNAcylated RNA Pol II. The development of the O-GlcNAc-specific antibodies has been challenging because glycoproteins are self-antigens, and the modification is small in size relative to the rest of the antigen [8]. To identify the localization of O-GlcNAcylated transcription machinery on chromatin, we developed a strategy to enrich for the O-GlcNAcylated chromatin where RNA Pol II operates (**Fig. 2A**). In this strategy, we first isolate O-GlcNAcylated chromatin using succinylated wheat-germ agglutinin (WGA), a plant-derived lectin that has high affinity to the O-GlcNAc modification [29]. We reasoned that it is possible to elute O-GlcNAcylated proteins from the WGA lectin using excessive GlcNAc, which competes with the binding. Indeed, this strategy revealed almost complete elution of the O-GlcNAcylated proteins from the lectin (**Fig. 2B**). After the elution, we performed traditional ChIP against RNA Pol II from the elute to selectively isolate for the O-GlcNAcylated chromatin that was associated with the polymerase. We termed this protocol as WGA re-ChIP (wChIP), and, for the simplicity, we refer to this purified fraction as ‘O-GlcNAcylated RNA Pol II’.

**Figure 2.**
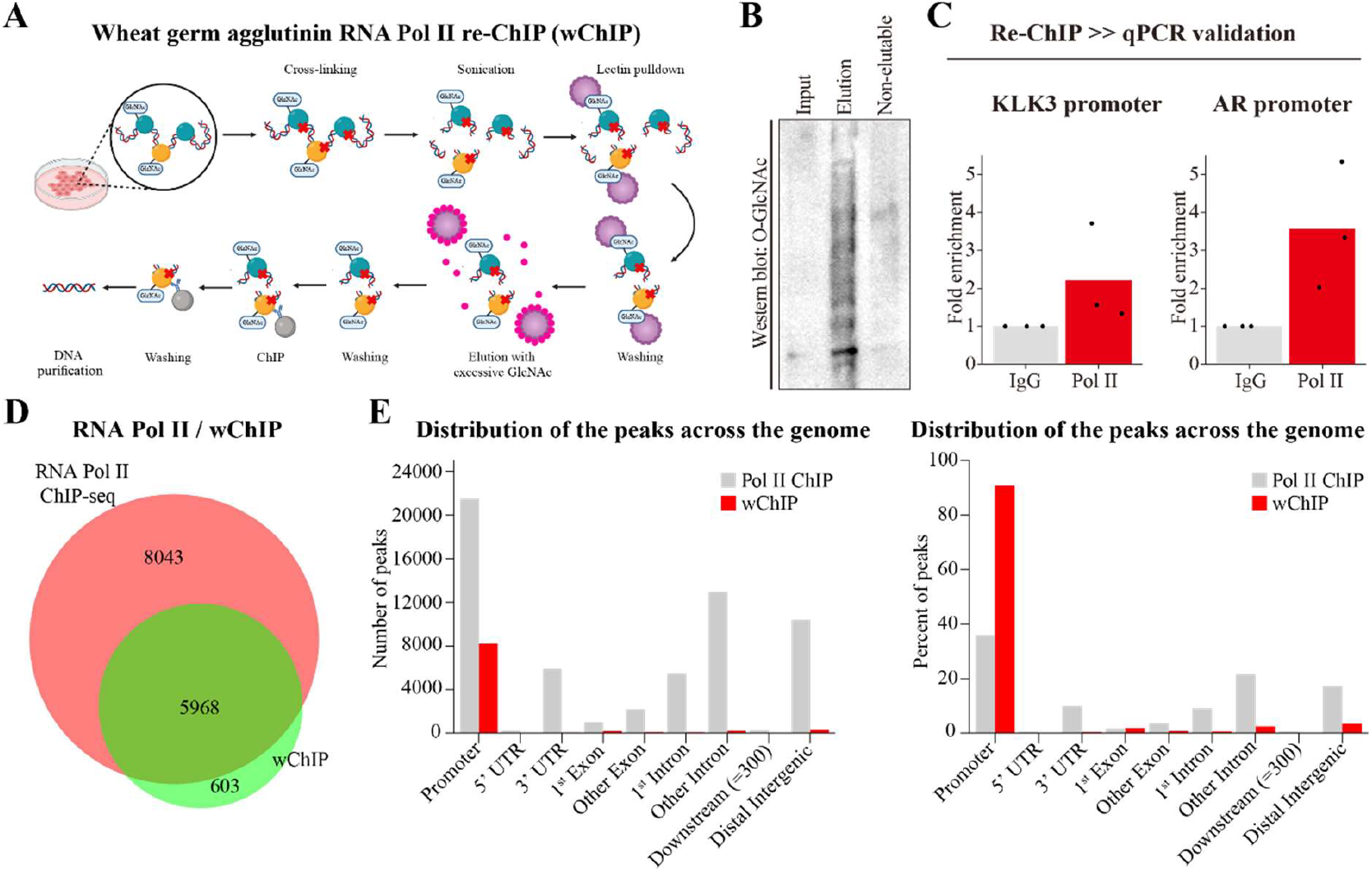
O-GlcNAcylated RNA Pol II localizes on promoters. **A)** Schematic of the WGA re-ChIP (wChIP) approach to identify chromatin binding sites of O-GlcNAcylated RNA Pol II. Briefly, chromatin-associated proteins are cross-linked to the DNA, DNA is fragmented with sonication and O-GlcNAcylated chromatin is isolated using wheat germ agglutinin (WGA), a lectin that has high affinity to GlcNAc. O-GlcNAcylated chromatin is eluted with excessive GlcNAc and the DNA bound by the O-GlcNAcylated RNA Pol II is isolated with traditional ChIP against the polymerase. Created with BioRender.com. **B)** Addition of GlcNAc successfully eluted almost all O-GlcNAcylated proteins from lectin. Western blot detection of O-GlcNAcylated proteins for samples collected at different stages of wChIP: sample collected before pulldown with WGA lectin (Input), proteins eluted from the lectin using excessive GlcNAc (Elution) and proteins that remained bound to the lectin (Non-elutable) (n: 1). **C)** 22RV1 cells were subjected to wChIP after which qPCR was used to assess presence of O-GlcNAcylated RNA Pol II on the promoters of genes encoding for androgen receptor (*AR*) and prostate-specific antigen (*KLK3*) (n: 3). **D)** Venn diagram of genes at which the O-GlcNAcylated RNA Pol II is located compared to genes at which RNA Pol II is located. Genes bound by O-GlcNAcylated RNA Pol II were identified with wChIP (binding sites common for three biological replicates). Chromatin binding sites of RNA Pol II were determined by re-analysis of ChIP-seq data reported in Takayama et al. [31] (GSE146886). Both datasets collected from 22RV1 cells. **E)** Number (left) and percentage (right) of wChIP and RNA Pol II ChIP-seq (Takayama et al. [1] (GSE146886)) binding sites at the given genomic regions.

To validate enrichment of O-GlcNAcylated RNA Pol II, we focused on the transcriptionally active promoters and analyzed our wChIP samples using qPCR. We selected two genes that are expressed in the 22RV1 cell line we use in our experiments (Kallikrein Related Peptidase 3: *KLK3* and Androgen Receptor: *AR*) [30]. Indeed, we found that the O-GlcNAcylated polymerase is enriched for promoter regions of these transcriptionally active genes (**Fig. 2C**).

Next, we sequenced the wChIP samples to get an unbiased view of the localization of the O-GlcNAcylated RNA Pol II on chromatin. This approach identified 9037 chromatin sites bound by the O-GlcNAcylated RNA Pol II that mapped to 6 571 genes (**Fig. 2D**). To relate localization of the O-GlcNAcylated RNA Pol II to the overall RNA Pol II, we re-analyzed a previously published RNA Pol II ChIP-seq dataset generated from the same cell line (GSE146886) [31] and compared the binding sites to our wChIP-seq results. As expected, we observed that most of the genes bound by the O-GlcNAcylated RNA Pol II (91 %) are also present in the traditional RNA Pol II ChIP-seq (**Fig. 2D**). Interestingly, the percentage of O-GlcNAcylated RNA Pol II on promoter regions was more than two-fold greater than for the total RNA Pol II (91 % and 36 % of the identified binding sites, respectively) (**Fig. 2E**). These data show that the O-GlcNAcylated RNA Pol II is predominantly located at the promoters. To summarize our major results so far, we developed wChIP method and show that the O-GlcNAcylated RNA Pol II is localized on the promoters. Next, we wanted to find out if O-GlcNAcylated transcription machinery is located on the transcriptionally active or silent genes.

### O-GlcNAcylated RNA Pol II localizes on promoters of transcriptionally active genes

We hypothesized that O-GlcNAc-mediated enrichment of RNA Pol II on promoters drives transcription of these genes. Besides driving the active transcription, polymerases can be poised on promoters of transcriptionally inactive genes to facilitate rapid activation of transcription at a later timepoint [32]. To probe if the O-GlcNAcylated chromatin where RNA Pol II is found at is transcriptionally active, we studied the enrichment of wChIP data on highly expressing genes. We identified the highly and lowly expressed genes from previously published RNA-seq dataset from the same cell line (GSE221263). For each of the three biological replicates in the RNA-seq data, we selected 500 genes with the highest expression based on reads per kilobase per million mapped reads (RPKM) values. For our panel of highly expressed genes, we selected those genes that were common for all the three biological replicates (486 out of 500). Similarly, we constructed a set of genes that are not expressed by selecting all genes where no RNA-seq reads were identified (RPKM = 0) in any of the biological replicates. This approach yielded a set of 2 867 genes with no expression based on RNA-seq. We then studied the enrichment of our wChIP reads at these highly and lowly expressed genes. We observed an almost 2-fold and 5-fold enrichment of O-GlcNacylated RNA Pol II on the transcription start sites of the highly expressed genes compared to all genes and the lowly expressed genes, respectively (**Fig 3A**). As expected, also the total RNA Pol II was enriched on the highly expressed genes (**Suppl. Fig. 2**). These results show that the O-GlcNAcylated RNA Pol II is associated with promoters of highly expressed genes.

**Figure 3.**
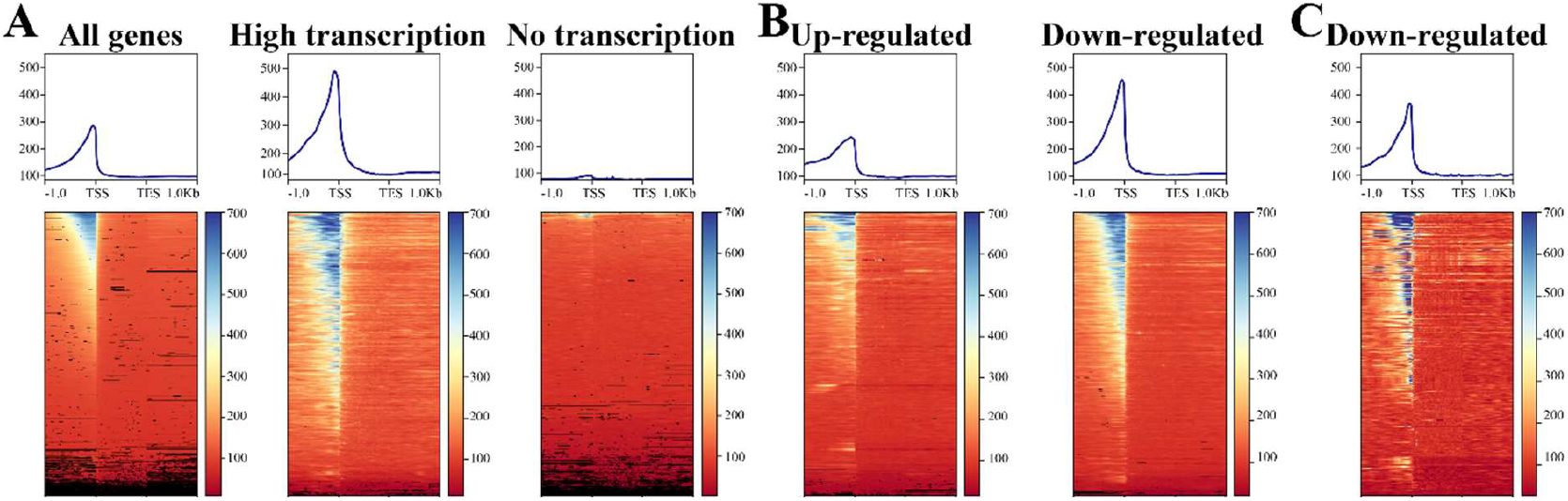
O-GlcNAcylated RNA Pol II is enriched on the highly transcribed genes. **A)** wChIP signal at protein-encoding genes that are highly expressed or lowly expressed. Highly and lowly expressed genes were determined by re-analysis of publicly available RNA-seq dataset (GSE221263). For each of the three biological replicates, 500 highest expressing genes were selected and common genes among these top 500 genes constitute the panel of highly expressed genes (in total 486 genes). Panel of lowly expressed genes consists of genes for which there was no identified reads (RPM = 0) in any of the biological replicates (in total 2 867 genes). **B)** wChIP signal at genes downregulated or upregulated by OGT inhibition (OSMI-2) in prostate cancer cells (LNCaP). Differentially expressed genes in response to OGT inhibition were previously reported (p.adj<0.05, upregulated (Log2FC>0): 699 genes and downregulated (Log2FC<0): 1139 genes) [9]. **C)** wChIP signal at genes differentially expressed in HEK cells in response to OGT inhibition with OSMI-2. Differentially expressed genes were identified from RNA-seq count matrices downloaded from the GEO database (GSE138783, p.adj<0.05 (Log2FC<0): downregulated: 74 genes) [33].

Next, we asked whether the transcription of genes harboring high levels of RNA Pol II O-GlcNAcylation are sensitive to OGT inhibition. We studied the coverage of wChIP reads on the genes whose expression was previously reported to be altered by OGT inhibition in prostate cancer cells [9]. Indeed, we observed that the O-GlcNAcylated RNA Pol II was clearly enriched at transcription start sites of the genes that were downregulated after OGT inhibition in contrast to the genes found to be upregulated after targeting OGT (**Fig. 3B**). To ascertain these results, we studied wChIP read coverage also at genes previously reported to be downregulated by OGT inhibition in human embryonic kidney (HEK) cells [33] and confirmed that the O-GlcNAcylated transcription machinery is enriched at transcription start sites of these genes (**Fig. 3C**). These data show that transcription of genes that are associated with O-GlcNAcylated RNA Pol II is sensitive to OGT inhibition.

We have so far shown that RNA Pol II O-GlcNAcylation position the polymerase on the promoters of the highly transcribed genes. We reasoned that RNA Pol II O-GlcNAcylation regulates its localization by recruiting a distinct set of proteins to the polymerase. However, we did not yet know which proteins bind to the O-GlcNAcylated RNA Pol II.

### Site-selective O-GlcNAcylation of RNA Pol II promotes association with KPNB1

We hypothesized that RNA Pol II O-GlcNAcylation regulates assembly of specific proteins to the polymerase. Since phosphorylation and O-GlcNAcylation have been reported to be mutually exclusive [13], we reasoned that depletion of RNA Pol II phosphorylation should augment the polymerase O-GlcNAcylation. To test this, we used inhibitors against the major transcriptional kinases (CDK7: 500 nM YKL-5-124 [34], CDK9: 20 nM NVP2 [35] and CDK12/13: 500 nM THZ531 [36]), and also depleted and augmented O-GlcNAcylation using compounds targeting OGT (20 μM OSMI-4 [37]) and OGA (20 μM Thiamet G [38]), respectively. The doses were selected based on our previous study [10]. We treated cells with these inhibitors for four hours after which we isolated O-GlcNAcylated proteins with WGA and detected the proteins of interest using western blot. Interestingly, inhibition of the transcription initiation kinase CDK7 increased O-GlcNAcylation of RNA Pol II, inhibition of CDK12/13 decreased it, while depleting CDK9 activity did not have a consistent effect (**Fig. 4A**). As expected, inhibition of OGT decreased RNA Pol II O-GlcNAcylation while OGA inhibition increased it (we note that in one of the four replicates the decrease of RNA Pol II O-GlcNAcylation for OGT inhibition is not seen). CDK7 predominantly phosphorylates RNA Pol II on Ser-5, and our lectin pulldown experiments therefore propose that OGT and CDK7 compete for the same site, Ser-5, for O-GlcNAcylation and phosphorylation.

**Figure 4.**
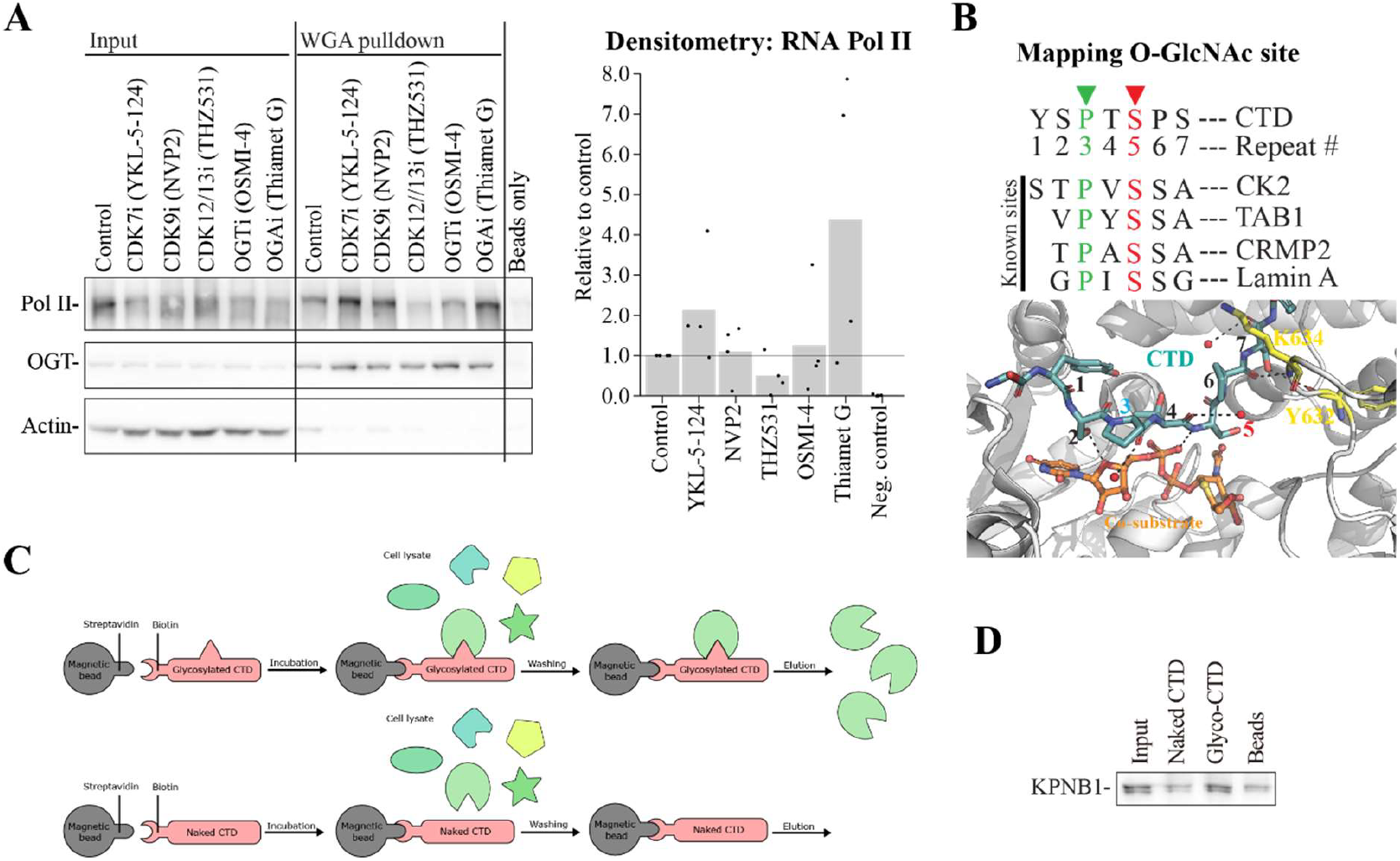
RNA Pol II carboxy-terminal domain O-GlcNAcylation at serine 5 promotes association with KPNB1. **A)** 22RV1 cells were treated with inhibitors of cyclin-dependent kinases CDK7 (500 nM YKL-5-124), CDK9 (20 nM NVP-2) and CDK12/13 (500 nM THZ531) as well as with inhibitors of OGT (20 μM OSMI-4) and OGA (20 μM Thiamet G) for four hours. Cell lysates were subjected to wheat-germ agglutinin (WGA) -lectin pulldown to isolate O-GlcNAcylated proteins and analyzed with western blot. Western blot signal was quantified with densitometry (n: 4). **B)** Sequence of consensus RNA Pol II CTD heptapeptide repeat. Proline amino acid precedes the O-GlcNAcylation site by one amino acid in the well-established OGT-substrates. When Ser-5 of the CTD is modeled (or hypothesized) as the O-GlcNAcylation site, the CTD could form extensive interactions with both the amino acids in the OGT active site and the OGT co-substrate (CTD-numbering as in CTD heptapeptide). Generated by modeling the CTD peptide into the OGT active site (based on the published structure 4N3B). **C)** Schematic of the pulldown of the RNA Pol II CTD binding partners. Briefly, biotinylated CTD peptides consisting of two repeats of the canonical heptapeptide repeat (Tyr1-Ser2-Pro3-Thr4-Ser5-Pro6-Ser7) either without post-translational modification (naked) or O-GlcNAcylated at Ser-5 were coupled to streptavidin-coated beads. Bead-coupled CTD peptides were incubated with 22RV1 cell lysate, and the binding partners were identified with mass spectrometry. **D)** Western blot validation of KPNB1 as a binding partner of CTD O-GlcNAcylated at Ser5 (n: 2).

We compared the consensus sequence of the CTD heptapeptide to the sequences of known OGT substrates (**Fig. 4C**). To our delight, the CTD sequence shares common features with OGT substrates [39]. In more detail, validated OGT-substrates casein kinase (CK2) [40], TAK1-Binding Protein 1 (TAB1) [41], Collapsin Response Mediator Protein-2 (CRMP2) [42], and Lamin A [43] all have proline in the -1-position preceding the O-GlcNAcylation site (**Fig. 4B**, top). Next, we manually modeled the CTD heptapeptide into the OGT’s active site based on a published structure (PDB id: 4N3B [44]). Modeling Ser-5 as the O-GlcNAcylation site could preserve the extensive interaction network within the active site (**Fig. 4B**, bottom), suggesting that Ser-5 is likely to be an O-GlcNAcylation site.

We next sought to identify the binding partners of RNA Pol II that is O-GlcNAcylated at Ser-5. We used chemically synthetized CTD peptides that were O-GlcNAcylated specifically at this site to pull down the binding partners from prostate cancer cell lysate and identified the proteins with mass spectrometry (**Fig. 4C**). Unmodified CTD peptide and the affinity matrix-only were used as controls. We validated karyopherin subunit beta 1 (KPNB1) to be associated more with the O-GlcNAcylated peptide than with the unmodified peptide or the affinity matrix (**Fig. 4D**).

To summarize our major findings so far, we found that RNA Pol II O-GlcNAcylation is dynamically regulated in response to CDK7 inhibition and identified CTD O-GlcNAcylation on Ser-5 as a regulator of interaction between KPNB1 and the polymerase. We note that the mass spectrometry-based identification was done only once, and it is therefore likely that there is a high number of false positive and / or negative hits: this is why we only focus on one protein that we successfully validated. Next, we wanted to understand the functional significance of this interaction.

### Combined inhibition of OGT and KPNB1 is toxic to prostate cancer cells

KPNB1 mediates the nuclear import of proteins [45], and we therefore hypothesized that O-GlcNAcylation of RNA Pol II is important for its nuclear import. To test this hypothesis, we performed fractionation experiments and detected RNA Pol II levels in different cellular compartments after treatment with inhibitors of OGT (OSMI-4 [37]) and KPNB1 (Importazole [46]). Indeed, combined inhibition of OGT and KPNB1 reduced nuclear and chromatin levels of RNA Pol II (**Fig. 5A**). We propose that the effects on chromatin-levels are secondary effects due to defective nuclear import. These data show that high activity of both OGT and KPNB1 are required to maintain high RNA Pol II levels in the nucleus and chromatin.

**Figure 5.**
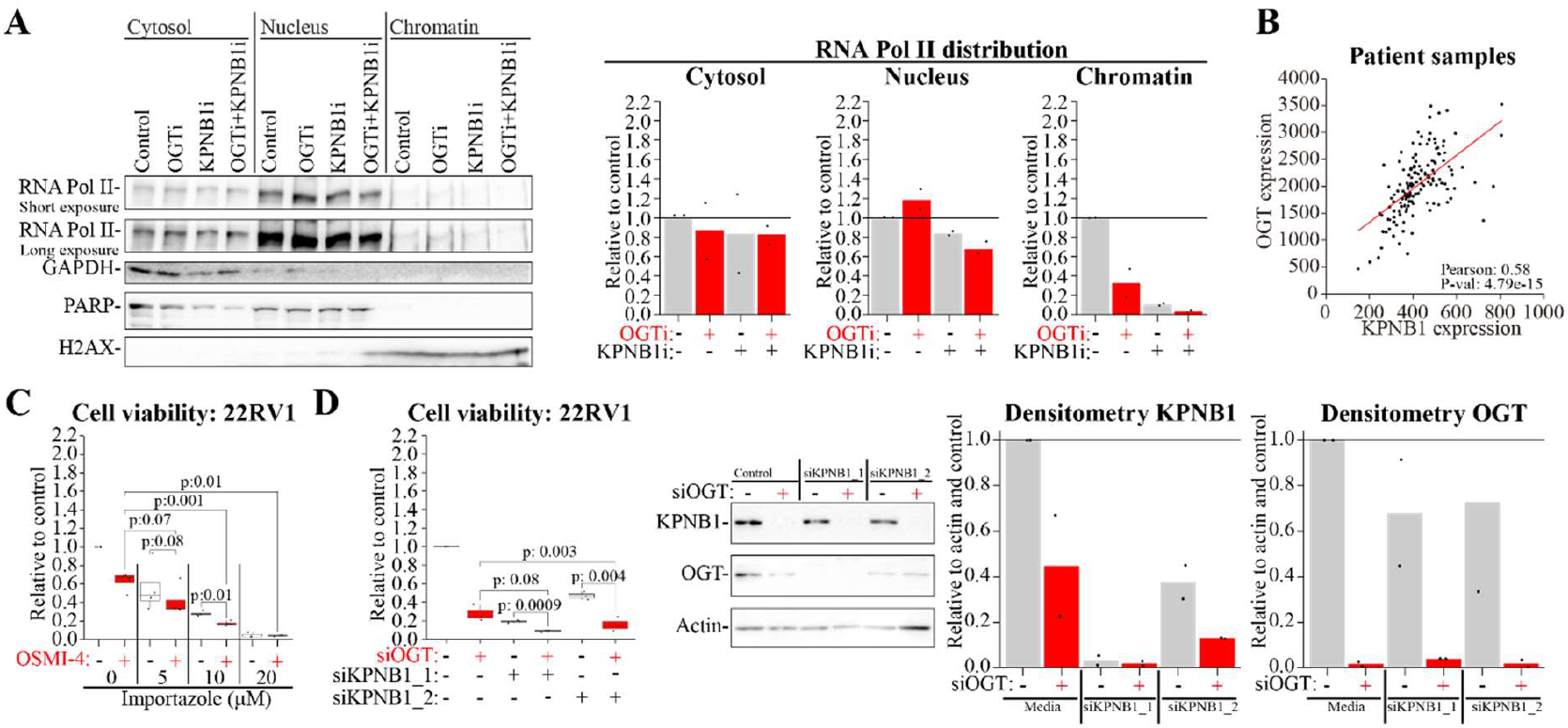
Inhibition of OGT and KPNB1 is combinatorically toxic to prostate cancer cells. **A)** 22RV1 cells were treated for 24h with OGT inhibitor OSMI-4 (20 μM) and/or KPNB1 inhibitor Importazole (20 μM) after which the fractionation was performed. Levels of RNA Pol II and loading controls for each fraction (cytoplasm: GAPDH, nuclear: PARP, chromatin: H2AX) were identified with western blotting. Densitometry was used to quantitate signal intensity and signal was normalized to loading control (n: 2). **B)** OGT and KPNB1 expression in prostate cancer patient samples (Data: Taylor et al. [69] accessed through betastasis.com). **C)** 22RV1 cells were treated with indicated doses of Importazole alone or in combination with OSMI-4 (20 μM) for four days. Cell viability was measured using CellTiterGlo (n: 3-4). **D)** Left: 22RV1 cell viability after knockdown of OGT and/or KPNB1. 22RV1 cells were reverse-transfected with indicated siRNAs for four days and cell viability was assessed with CellTiterGlo (n: 3). Right: western blot to confirm the knockdown of the target proteins. Cells were lysed three days after the reverse-transfection, the levels of OGT and KPNB1 was assessed with western blot and the signal quantified with densitometry (n: 2). C and D: Paired two-tailed Student’s t-test was used to assess the significance.

We reasoned that OGT and KPNB1 are important for cancer cells that rely on high levels of transcription to maintain their proliferation. Previously, both OGT and KPNB1 have been reported to be over-expressed in prostate cancer [19, 47]. Indeed, we found a significant positive correlation between the expression levels of OGT and KPNB1 in patient tumours (**Fig. 5B**). These data propose that prostate cancer cells benefit from high activity of OGT and KPNB1.

To directly assess if prostate cancer cells are addicted to high activity of OGT and KPNB1, we used proliferation assays. Indeed, combinatorial targeting of OGT and KPNB1 significantly decreased viability and reduced colony-formation ability of prostate cancer cells (**Fig. 5C and Suppl. Fig. 3**). In contrast, cell line derived from normal prostate tissue (PNT1) was less sensitive to this combination (**Suppl. Fig. 3A**), which supports our proposal that cancer cells are more dependent on both OGT and KPNB1. To validate the efficacy of the combination treatment, we used knockdown experiments in prostate cancer cells. This revealed significant further decrease in cell viability for the dual knockdown when compared to single knockdowns (**Fig. 5D**). These experiments additionally revealed that knockdown of either OGT or KPNB1 reciprocally reduces each other’s levels, which further supports functional importance of high activity of both. To probe the potential translational value of our finding, we used ivermectin, a compound that targets KPNB1 but also KPNA [48] and is currently in clinical trials for metastatic triple negative breast cancer (clinicaltrials.gov identifier: NCT05318469). Indeed, combining OGT inhibitor OSMI-4 with Ivermectin drastically suppressed proliferation of prostate cancer cells but was less toxic to PNT1 cells (**Suppl. Fig. 4**). These data show that prostate cancer cells are addicted to the high activity of both OGT and KPNB1.

To summarize the major findings of our project, we propose that RNA Pol II O-GlcNAcylation promotes nuclear entry and positions the polymerase on the highly transcribed genes. We identified KPNB1 as a selective binding partner of the O-GlcNAcylated RNA Pol II and showed that combined inhibition of OGT and KPNB1 is toxic to prostate cancer cells. OGT inhibitors have never been tested in clinical trials, and it would be of interest to assess if inhibitors of transcriptional kinases can sensitize cells to compounds targeting KPNB1. Finally, there is significant value in developing tools that enable enrichment of the selectively O-GlcNAcylated RNA Pol II and mapping the interactome of the differentially O-GlcNAcylated polymerase – these should be established in the future. We propose that O-GlcNAcylation of RNA Pol II on Ser-5 positions the polymerase on the highly transcribed genes, and cancer cells hijack this to promote transcription.

## Materials and Methods

### Cell culture, compounds, and proliferation assays

22RV1 and C4-2 cell lines were acquired from the American Tissue Culture Collection (ATCC), and PNT1 cells were acquired from Sigma. 22RV1, C4-2 and PNT1 cells were maintained in the RPMI medium supplemented with 10% fetal bovine serum (FBS). All cells were allowed to adhere for at least one day prior to the treatment with inhibitors. Importazole [46], Ivermectin [48], NVP2 [35], OSMI-4 [37], Thiamet G [38], THZ531 [36], and YKL-5-124 [34] were obtained from MedChemExpress. To assess cell viability, we used CellTiter-Glo® 2.0 assay (Promega) according to manufacturer’s instructions. Crystal violet staining was performed as previously reported [49].

### Preparation of cell lysates, western blot-detection, lectin pulldown, and knockdown experiments

Cell lysates were prepared as previously described [19] with minor changes. All steps were performed at 4ºC unless otherwise stated. Briefly, cells were first washed with PBS, and then incubated in the cell lysis buffer (RIPA buffer: 10 mmol/L Tris-HCl, pH 8.0, 1 mmol/L EDTA, 0.5 mmol/L EGTA, 1% Triton X-100, 0.1% sodium-deoxycholate, 0.1% SDS, 140 mmol/L NaCl) supplemented with inhibitors of protease, phosphatase and O-GlcNAcase (Thiamet G) for 30 minutes. Then, the samples were centrifuged at 15 000 rpm for five minutes and the supernatant was collected. For western blot detection, protein concentration was determined with bicinchoninic acid (Pierce™ BCA Protein Assay Kit, Thermo Fisher Scientific). Antibodies used were from Santa Cruz Biotechnology: RNA Pol II (sc-47701, sc-56767), GAPDH (sc-47724), H2AX (sc-517336); from Abcam: O-GlcNAc (ab2739), Actin (ab49900), from Cell Signaling Technologies: OGT (24083), PARP (9532); and from ABclonal: KPNB1 (A8610). Western blot signal intensity was quantified using Image Lab version 6.0 (Bio-Rad). Lectin pulldown was performed as previously described [50]. In knockdown experiments, RNAiMax (Thermo Fisher Scientific) was used to transfect the Silencer® Select siRNAs against OGT (s16094) and KPNB1 (s7918 and s7919). Knockdown was performed for four days, and the cell viability was assessed with CellTiter-Glo 2.0 assay (Promega). For the validation of knockdown, partial medium change was performed after 2 days, and cell lysate was collected after 3 days.

### Cell fractionation

Cell fractionation was performed according to previously reported protocol [51] with minor adjustments. Briefly (all steps at 4ºC), cell pellets were lysed with 200 µL NETN100 buffer (50 mM Tris-HCl, pH 7.4, 2 mM EDTA and 100 mM NaCl) for 30 minutes. After centrifugation at 3 000 rpm for 10 minutes, supernatants were collected (cytoplasmic fraction). Insoluble pellets were washed twice with PBS and incubated with 200 µL NETN300 buffer (50 mM Tris-HCl, pH 7.4, 2 mM EDTA and 300 mM NaCl) for 30 minutes. After pelleting with centrifugation at 3 000 rpm for 10 minutes, supernatant was collected (nuclear fraction). Insoluble pellet was washed twice with PBS, centrifuged at 3 000 rpm for 10 minutes and treated with 100 µL 0.2N HCl for 10 minutes. Finally, the sample was centrifuged at 15000 rpm for 10 minutes and the supernatant was neutralized with 25 µL 1N NaOH (chromatin fraction).

### WGA re-ChIP (wChIP)

Parts of WGA ChIP (wChIP) were performed as previously reported in [52] with minor adjustments. Chromatin-associated proteins were cross-linked to the chromatin with 1% formaldehyde for 10 minutes at room temperature prior to quenching with 125 mM glycine for 10 minutes. The following steps were performed at 4ºC unless indicated otherwise. After harvesting the cells and washing them with PBS, nuclear lysates were isolated. First, cells were incubated with 5 ml lysis buffer 1 (50 mM Hepes-KOH, 140 mM NaCl, 1 mM EDTA, 10% glycerol, 0.3% Triton X-100) for 10 minutes. Then, nuclei were pelleted by centrifuging with 2000 rpm for 5 minutes. After incubating the nuclei for 5 minutes in the 5 ml of lysis buffer 2 (10 mM Tris-HCl pH8.0, 200 mM NaCl, 1 mM EDTA, 0.5 mM EGTA) and centrifuging at 2000 rpm for 5 minutes, 3 ml lysis buffer 3 (10 mM Tris-HCl pH8.0, 100 mM NaCl, 1 mM EDTA, 0.5 mM EGTA, 0.1% Sodium Deoxycholate) was added and samples were divided into fractions of volume of 270 μl. Samples were sonicated in water bath sonicator (Bioruptor) for 13 cycles where each cycle consisted of 30 seconds of sonication and 30 seconds of rest. After sonication, 30 μl 10% of Triton X-100 was added, sample was centrifuged at 15 000 rpm for 10 minutes and supernatant was collected.

Next, O-GlcNAcylated protein-chromatin complexes were isolated. Agarose beads coated with succinylated WGA (VectorLabs, AL-1023S-5) were washed three times with lectin washing buffer (0.1 % Tween, 150 mM NaCl, 10 mM Tris–HCl pH 8.0). Samples were incubated with the beads for four hours with agitation after which the beads were washed three times with the lectin washing buffer. O-GlcNAcylated protein-chromatin complexes were eluted with 150 μl 500 mM glucosamine for 30 minutes at room temperature with agitation. Samples were centrifuged at 3000 rpm for 3 minutes. Then the supernatant (150 μl) was diluted to 1400 μl 0.5% BSA in lysis buffer 3 supplemented with protease inhibitor and 50 μM O-GlcNAcase inhibitor (Thiamet G).

Protein A/G magnetic beads (MedChemExpress, HY-K0202) were first washed twice with 0.5% Tween in PBS. Next, the beads were incubated with 10 μg of antibodies (RNA Pol II: sc-47701 from Santa Cruz Biotechnology, IgG: 015-000-003 from Jackson ImmunoResearch) and protease inhibitor in 0.5% BSA in PBS for 8 hours in +4ºC with agitation. Antibody-bead complexes were washed four times with 0.5% BSA in PBS, after which samples were incubated with beads overnight. The next day, beads were washed five times with LiCl washing buffer (50 mM Hepes-KOH pH 7.6, 500 mM LiCl, 1 mM EDTA, 1% Triton X, 0.7% Sodium Deoxycholate) and once with TE buffer (1 mM EDTA pH 8.0, 10 mM Tris-HCl). Samples were eluted in elution buffer (50 mM Tris-HCL pH 8.0, 10mM EDTA, 1% SDS) and decrosslinking was performed overnight at 65ºC.

RNA and proteins were degraded by incubating the sample first with 80 μg of DNAse-free RNAse A (Thermo Fisher Scientific, EN0531) for 1 hour at 37ºC and then with 80 µg Proteinase K (Thermo Fisher Scientific, EO0491) for 1 hour at 55ºC. DNA was isolated first with phenol/chloroform/isoamyl alcohol (25:24:1 Thermo Fisher Scientific, BP17521-100) extraction and then with chloroform extraction. The DNA was precipitated by adding 0.13 volumes of 3M Sodium Acetate and three volumes of 100% ethanol after which the sample was stored in -80°C for at least 2 hours. Finally, sample was air-dried and resuspended in nuclease-free water. Following primers were used in qPCR: androgen receptor promoter (F-TCGACCATTTCTGACAACGC, R-GAAGCTGTTCCCCTGGACTC), and KLK3 promoter (F-CCTAGATGAAGTCTCCATGAGCTACA, R-GGGAGGGAGAGCTAGCACTTG). Sequencing and library preparation of wChIP samples was purchased from the DNA Sequencing and Genomics Laboratory (BIDGEN, Helsinki, Finland).

### CTD-biotin pulldown and mass spectrometry

Chemically synthetized biotin conjugated CTD peptides consisting of two repeats of heptapeptide tyrosine-serine-proline-threonine-serine-proline-serine were obtained from JPT Peptide Technologies. Two types of peptides were used: peptides O-GlcNAcylated at both serine 5 residues and peptides without post-translational modifications (naked). For the biotin-CTD pulldown, cell lysates were incubated in lysis buffer (0.5% TritonX, 150 mM NaCl, 50 mM Tris-HCl pH 7.5, supplemented with 25 μM Thiamet G, 10 μM OSMI-1, protease inhibitor and phosphatase inhibitor) for 15 minutes and sonicated in water bath sonicator (BioruptorPlus, Diagenode) for 25 minutes with cycle: 30 sec on, 30 sec rest. After washing streptavidin-coated magnetic beads (Promega, Z548C) three times with PBS, CTD peptides were incubated with beads in the presence of protease inhibitor for two hours at the room temperature with agitation. Next, beads were washed three times with PBS and incubated with the cell lysate for four hours at the room temperature with agitation. Finally, the beads were washed three times with PBS. Mass spectrometry analysis of samples was performed by the University of Helsinki Meilahti Clinical Proteomics Core Facility.

### Bioinformatics

ChIP-seq and wChIP-seq results were analyzed in the following manner. First, the raw fastq files were subjected to quality control using FastQC [53] after which they were aligned to human genome Hg38 using Bowtie2 aligner [54]. Second, Samtools [55] was used to convert the resulting Sequence Alignment Map (SAM) files to binary format (BAM) and sorted. Third, MACS2 [56] was used to call peaks and Bedtools [57] to find binding sites common for all the replicates. Finally, ChIP-seq signal at the selected genome regions was calculated and plotted using deepTools [58].

Pan-cell line list of the binding sites of O-GlcNAcylated chromatin was constructed using previously published O-GlcNAc ChIP-seq and click chemistry-based COGC-seq datasets. Files containing the ChIP peaks were downloaded from the GEO database: colon cancer (HT-2) (GSE93656) [26], human embryonic kidney (HEK293) (GSE51673) [27] and Burkitt lymphoma (BJAB) (GSE86154) [28]. Binding sites for prostate cancer cell lines (LNCaP and PC3) were used as reported in a previous study [15] and breast cancer (MCF-7) [16] data was obtained from the article supplement. HEK293 dataset (GSE36620) [14] was re-analyzed starting from fastq files. Genome annotations were lifted to hg38 assembly using UCSC tool liftOver [59], and Bedtools [57] was used to find binding sites common for all the datasets.

To identify genes lowly and highly expressed in 22RV1 cell line, RNA-seq data was downloaded from the GEO database (GSE221263). After quality control with FastQC [53], reads were aligned to human genome Hg38 with STAR aligner [60]. The resulting SAM files were converted to BAM format and sorted using Samtools [55]. Reads were counted with featureCounts [61]. Reads Per Kilobase Million (RPKM) normalization was performed with edgeR [62].

Genes differentially expressed after OGT inhibition in HEK cells were identified from RNA-seq count matrices downloaded from the GEO database (GSE138783) [33]. Differentially expressed genes were identified with DESeq2 [63]. The differentially expressed genes after OGT inhibitor were used from our previous publication [9]. Genes with adjusted p-value less than 0.05 were considered significant.

We annotated the identified binding sites using R package ChIPseeker [64]. Venn diagrams were plotted using BioVenn [65] and RStudio. Enrichment analyses were performed with Enrichr (KEGG 2021 Human pathways) [66-68]. For statistical analysis, paired two-tailed Student’s t-test was used unless otherwise mentioned.

## Author contributions

SP performed most of the experiments as well as bioinformatics and statistical analyses and plotted figures and wrote the manuscript together with HMI. XL performed computational docking analysis and plotted the associated figures. AG participated in the development of CTD pulldown protocol and acquired resources for mass spectrometry. HMI conceptualized the study, supervised the project, participated in wChIP experiments and obtained resources.

## Funding

SP is supported by the Emil Aaltonen Foundation, Waldemar von Frenckells Stiftelse, Otto A. Malm Foundation and University of Helsinki Doctoral Programme in Biomedicine. HMI is grateful for the funding from the Academy of Finland (Decision nr. 331324, nr. 335902 and nr. 358112), the Jenny and Antti Wihuri Foundation, the Finnish Cancer Foundation, and Sigrid Juselius Foundation.

## Acknowledgements

We wish to acknowledge CSC – IT Center for Science, Finland, for the computational resources. Proteomic analyses were performed at the Meilahti Clinical Proteomics Core Facility. Therefore, we acknowledge the facility and its supporters: HiLIFE and Biocenter Finland. We wish to also express our gratitude to the DNA Sequencing and Genomics Laboratory (BIDGEN, Helsinki, Finland) where the library preparation and sequencing of the samples was performed.

## Conflict of interest

Authors have no conflict of interest to declare.

## Notes

### Competing Interest Statement

The authors have declared no competing interest.

