## Supplementary figures and supplementary figure legends for "RNA polymerase II O-GlcNAcylation promotes nuclear entry to drive transcription"

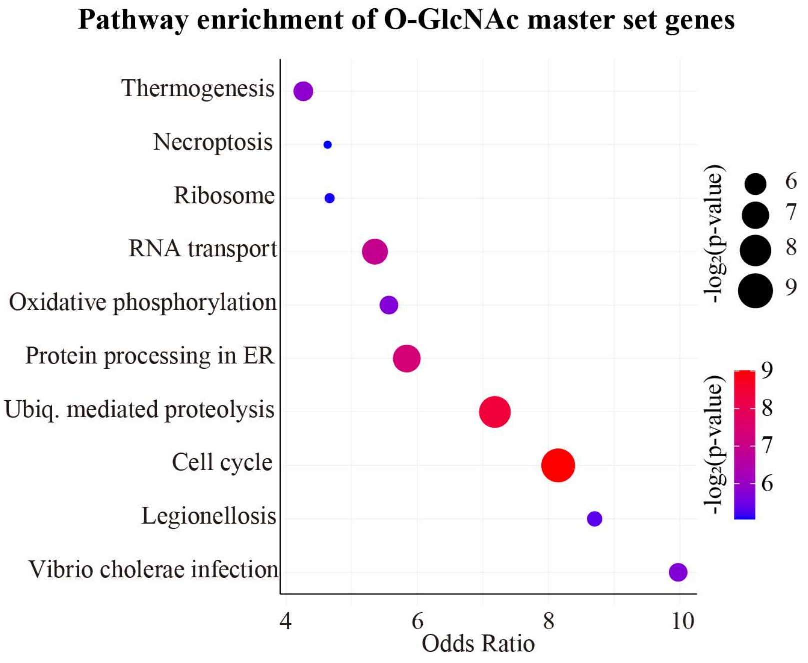


**Supplementary figure 1. KEGG enrichment analysis of genes at which there was an overlap of O-GlcNAcylated chromatin regions between all the seven datasets.** Enrichment analysis performed with Enrichr using KEGG 2021 Human pathways [66-68]. None of the gene sets were significant based on the p-value adjusted with Benjamin-Hochberg method.

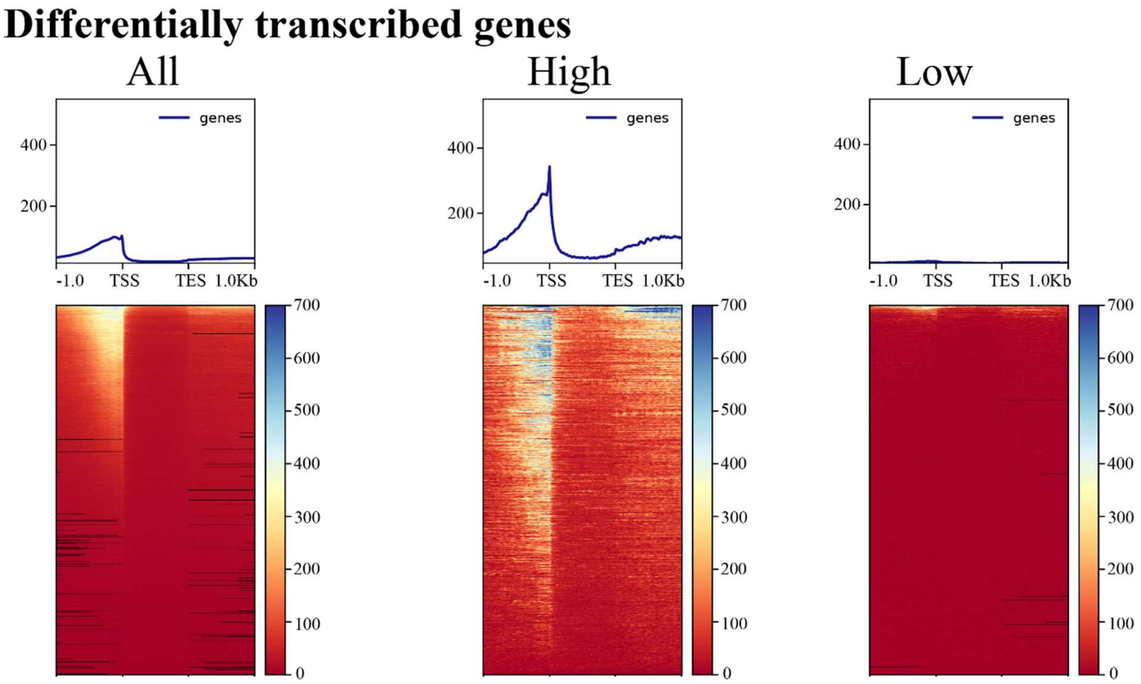


**Supplementary figure 2. RNA Pol II is localized on promoters of highly expressed genes.** RNA Pol II ChIP-seq signal at protein-encoding genes that are highly or lowly expressed. Panels of the genes that are highly or lowly expressed were established through re-analysis of publicly available RNA-seq dataset (GSE221263). For the panel of highly expressed genes 500 highest expressing genes were selected for each replicate and common genes among the replicates were selected (in total 486 genes). For the panel of lowly expressed genes, the genes for which there was not identified reads (RPM = 0) in any of the biological replicates were selected (in total 2 867 genes). Re-analysis of RNA Pol II ChIP-seq signal downloaded from GEO database (GSE146886) [31].

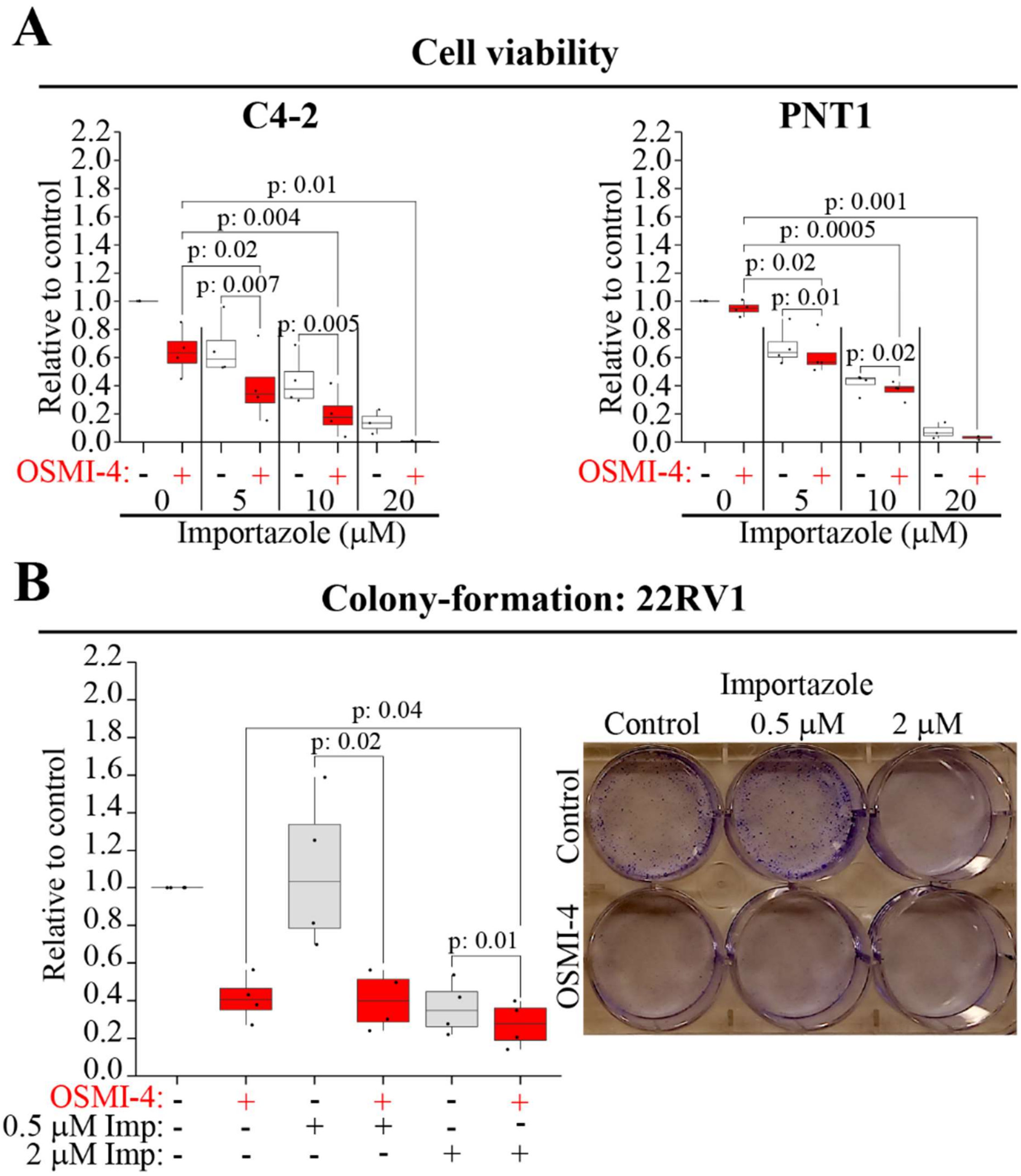


**Supplementary figure 3. Simultaneous inhibition of OGT and KPNB1 is toxic to prostate cancer cells. A)** Cell viability of CRPC cells (C4-2) and cells derived from normal prostate epithelia (PNT1) after treatment with indicated doses of KPNB1 inhibitor (Importazole) and/or OGT inhibitor (20 μM OSMI-4) for four days. Cell viability was assessed using CellTiterGlo. Significance was determined using paired two-tailed Student's t-test (n: 3-4). **B)** Colony-formation of 22RV1 cells after seven days of treatment with OSMI-4 (10 μM) and/or Importazole (0.5 μM or 2 μM) (n: 4). Statistical significance was assessed with paired two-tailed Student's t-test.

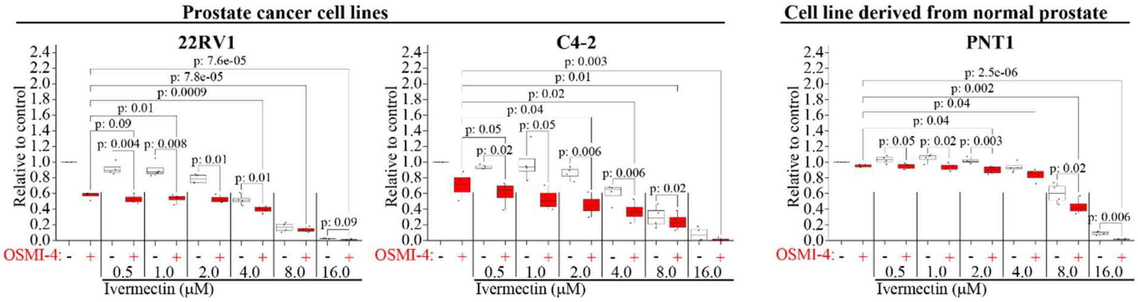


**Supplementary figure 4. Combined inhibition of OGT and KPNA/KPNB1 reduces viability of prostate cancer cells.** Viability of castration-resistant prostate cancer cells (22RV1 and C4-2), and a cell line representing normal prostate tissue (PNT1). Cells were treated for four days with indicated doses of KPNA/KPNB1 inhibitor (Ivermectin) and/or OGT inhibitor (20 μM OSMI-4) and cell viability was measured with CellTiterGlo. Statistical significance was assessed with paired two-tailed Student's t-test (n: 4).
